# Identification of dendritic cell-T cell interactions driving immune responses to food

**DOI:** 10.1101/2022.10.26.513772

**Authors:** Maria C.C. Canesso, Tiago B.R. Castro, Sandra Nakandakari-Higa, Ainsley Lockhart, Daria Esterházy, Bernardo S. Reis, Gabriel D. Victora, Daniel Mucida

## Abstract

The intestinal immune system must concomitantly tolerate food and commensals and protect against pathogens. Dendritic cells (DCs) orchestrate these immune responses by presenting luminal antigens and inducing functional differentiation of CD4^+^ T cells into regulatory (pTreg) or pro-inflammatory (Th) subsets. However, the exact nature of the DCs inducing tolerance or inflammation to dietary antigens has been difficult to define. Using an intestine-adapted Labeling Immune Partnerships by SorTagging Intercellular Contacts (LIPSTIC) combined with single-cell transcriptomics, we characterized DCs presenting dietary antigens in the context of tolerance or infection. At steady-state, migratory cDC1 and cDC2 DCs, but not resident DCs, were found to present dietary antigen to cognate CD4^+^ T cells. Whereas cDC2s promoted T cell activation, only cDC1s induced their differentiation into pTregs. Infection with the helminth *Strongyloides venezuelensis* abrogated cDC1 presentation of dietary antigens, preventing pTreg and oral tolerance induction. In contrast, *Heligmosomoides polygyrus* infection only partially affected cDC1s, allowing oral tolerance to be maintained. An expanded population of cDC2s that induced type-2 immunity during both helminth infections did not present dietary antigens, demonstrating that compartmentalized presentation of luminal antigens can prevent food-specific Th2 responses during inflammatory conditions. Our data uncover novel cellular mechanisms by which tolerance to food is induced and can be disrupted during infections.

## Introduction

The balance between pro-inflammatory and tolerogenic immune responses depend on interactions between dendritic cells (DCs) and T cells^1,2^. In the intestine, DCs take up food and microbial antigens from the intestinal lumen and then migrate to the gut-draining lymph nodes (gLNs) where they present these antigens to naïve T cells^3^. By inducing differentiation of regulatory or pro-inflammatory CD4^+^ T cell functional subsets during antigen presentation, DCs orchestrate finely-tuned immune responses to different intestinal antigens^4–7^. Although intestinal DC subsets with different tolerogenic and immunogenic functions have been described^8–11^, it remains unclear how DC-T cell interactions are organized to enable distinct immune responses given the density and diversity of luminal antigens constantly present in the gLNs. Furthermore, how diet or enteric infections regulate DC-T cell interactions to modulate immune outcomes is incompletely understood. Here we investigated the cellular mechanisms underlying the decision between tolerance or immunity to dietary antigens.

Oral tolerance, a process dependent on peripheral regulatory T cells (pTregs), is critical for preventing inflammatory responses to food antigens^12^. Migratory classical DCs are subdivided into two subsets, cDC1 (CD103^+^ CD11b^−^) and cDC2 (CD103^+/−^ CD11b^+^), both of which have been implicated in the induction of tolerogenic pTregs. Early studies examining CD103^+^ DCs documented their ability to induce pTregs in a manner dependent on TGF-β and the retinoic acid-producing enzyme retinaldehyde dehydrogenase (RALDH)^8,10,13,14^. However, oral tolerance is maintained upon genetic depletion of cDC1s, even though food antigen-specific pTreg induction is reduced^5^. Targeting cDC2 antigen presentation also reduces pTreg differentiation^15,16^, suggesting a potential compensatory mechanism among DC populations in promoting tolerance^5,17^. Full dissection of the role of DCs in oral tolerance has thus been limited by reliance on traditional knockout strategies, which have incomplete targeting efficiency and specificity and suffer from compensatory effects^1^.

## Results

Conclusive determination of which DCs dictate tolerance or inflammation in the gut requires the ability to precisely identify DCs presenting antigen in these contexts. To this end, we used an intestine-adapted version of LIPSTIC, which allows proximity-dependent labeling of intestinal cell-cell interactions *in vivo*^18^. LIPSTIC tracks the exact DCs that interact with naïve CD4^+^ T cells in an antigen-dependent manner, avoiding confounding factors due to parallel immune responses to other luminal stimuli. In the LIPSTIC system, CD40L (expressed by recently activated T cells^19^) and CD40 (expressed by DCs) are functionalized by fusion to Sortase A (SrtA) and its receptor G5, respectively. Injection of labeled SrtA substrate (the peptide LPETG conjugated to biotin) leads to its capture by donor SrtA-expressing T cells. DC-T cell interaction results in covalent transfer of substrate from T cell to DC^18^. Interacting DCs are thus detectable by flow cytometry with biotin-specific reagents (**Fig. 1a**).

**Figure 1.**
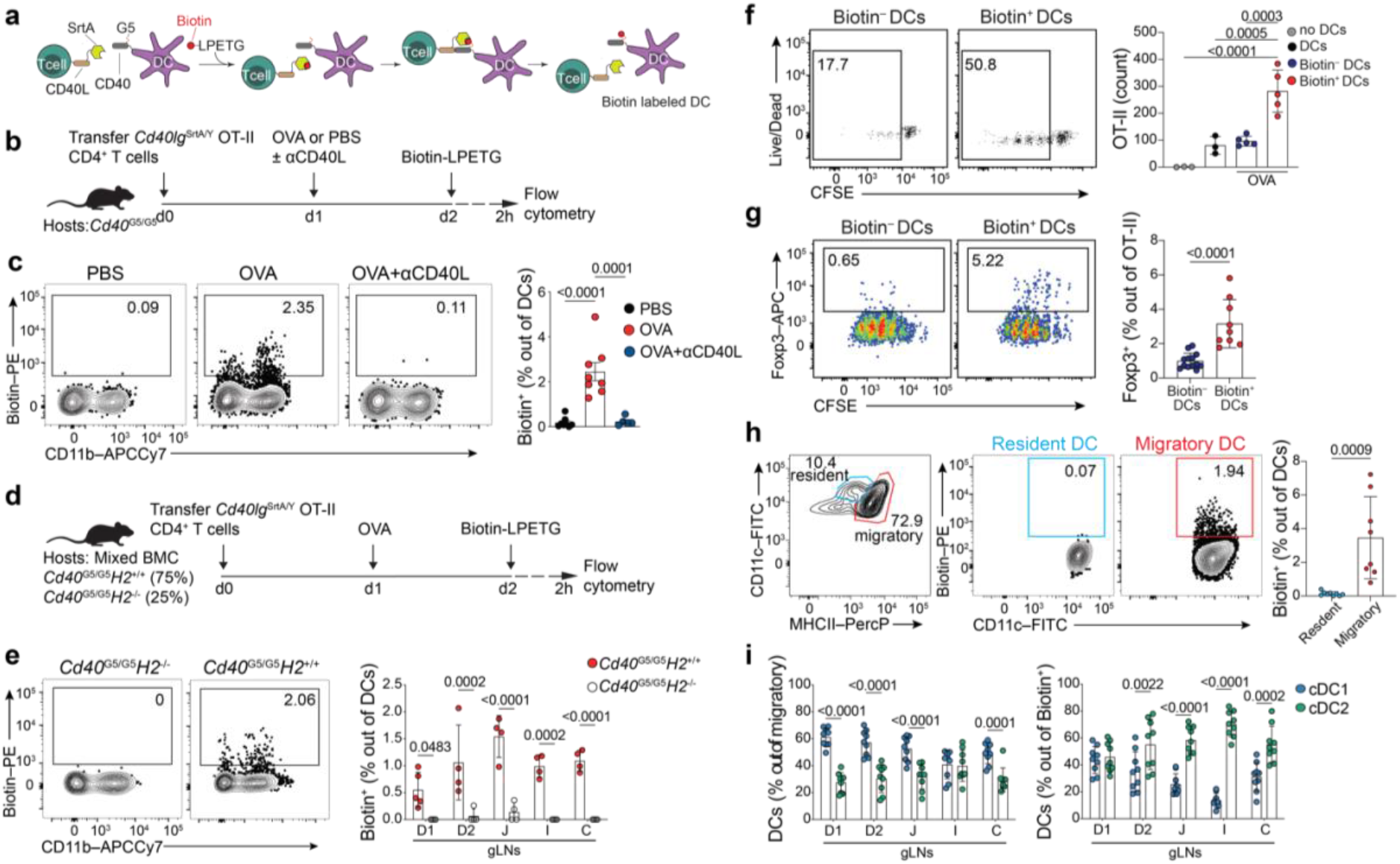
Using LIPSTIC to identify DCs presenting dietary antigens in the gLNs. **(a)** Schematic representation of LIPSTIC labeling of inter-cellular contacts *in vivo*. **(b, c, f-i)** CD45.2 *Cd40*^G5/G5^ mice were adoptively transferred with 1 × 10^6^ naive CD45.1 CD4^+^*Cd40lg*^SrtA/Y^ OT-II T cells prior to one dose of intragastric PBS, OVA or OVA + anti CD40L antibody. Cell-cell interaction was revealed by LIPSTIC protocol 24 h later. **(b)** Experimental setup for panel (c). **(c)** Representative flow plots showing percentage of labeled DCs in the duodenal gLNs (D-gLNs) (left) and quantification of data (right) (n = 3, 4 mice per group, pool of two independent experiments). **(d, e)** Mixed bone marrow chimera (BMC) mice reconstituted with *Cd40*^G5/G5^ and *Cd40*^G5/G5^;*H2*^−/−^ cells. Mice were adoptively transferred with 1 × 10^6^ naive CD4^+^*Cd40lg*^SrtA/Y^ OT-II T cells prior to one dose of intragastric OVA. Cell-cell interaction was revealed by LIPSTIC protocol 24 h later. **(d)** Experimental setup for panel (e). **(e)** Representative flow plots showing percentage of labeled DCs (n = 4 mice per group, representative of two independent experiments). **(f,g)** Sorted D-gLNs biotin^−^ or biotin^+^ DCs or DCs derived from OVA-naive mice were co-culture *in vitro* with naïve OT-II CFSE-labeled T cells for 96 h prior analysis. **(f)** Representative flow plots showing proliferation of OT-II T cells co-cultured with biotin^−^ or biotin^+^ DCs (left) and quantification of OT-II T cells per well at the end of the culture period with (right). Each dot represents one mouse (n = 3 to 5 mice per group, representative of two independent experiments). **(g)** Representative flow plots showing percentage of Foxp3^+^ cells among proliferated OT-II T cells co-culture with biotin^−^ or biotin^+^ DCs in the presence of exogenous OT-II peptide (left), and quantification of data (right). Each dot represents one mouse (n = 3, 4 mice per group, pool of three independent experiments). **(h, i)** CD45.2 *Cd40^G5/G5^* mice that were adoptively transferred with 1 × 10^6^ naive CD45.1 CD4^+^*Cd40lg*^SrtA/Y^ OT-II T cells prior to one dose of intragastric OVA. Analyses were carried out 24 h later. (**h**) Representative flow plots showing gating on resident and migratory DCs in D-gLNs (left), percentage of labeled resident and migratory DCs in D-gLNs (center), and quantification of data (right). Each dot represents one mouse (n = 3 mice per group, pool of three independent experiments). **(i)** Percentage of cDC1 and cDC2 out of total migratory DCs (left) and percentage of cDC1 and cDC2 out of biotin^+^ DCs (right). Each dot represents one mouse (n = 3 mice per group, pool of three independent experiments). D1, portal lymph nodes; D2, distal duodenum gLNs; J, jejunum; I, ileum; C, colon. *P*-values were calculated by unpaired *t*-test, one-way or two-way ANOVA.

To identify the intestinal DCs that present dietary antigens and prime cognate naïve CD4^+^ T cells, we adoptively transferred 1 × 10^6^ naïve ovalbumin (OVA)-specific CD40L-SrtA OT-II (*Cd40/g*^SrtA/Y^ OT-II) T cells into G5-CD40-expressing (*Cd40*^G5/G5^) mice. To induce a pTreg-dependent tolerogenic response, recipient mice received a single intragastric (i.g.) dose of PBS (control) or OVA 18h after cell transfer^3^. LIPSTIC labeling of interacting DCs in the gLNs was carried out by intraperitoneal (i.p.) injections of biotinylated LPETG substrate 22-24 h after i.g. OVA (**Fig. 1b**), a point at which CD40L is upregulated on the transferred OT-II cells (**Suppl. Fig. 1a, b**). LIPSTIC labeling was observed in a small fraction (approximately 2 to 3%) of total DCs in the gLNs of mice that received OVA, but not in control mice that received i.g. PBS (**Fig. 1c**). Increasing the number of transferred OT-II T cells did not increase DC labeling frequency, indicating that most if not all DCs presenting to OT-II T cells are captured (**Suppl. Fig. 1c**). Furthermore, DC labeling was fully inhibited by injection of a CD40L-blocking antibody (clone MR1) 22 h hours prior to substrate injection, confirming that CD40-CD40L interaction is required for labeling to take place (**Fig. 1c**). To evaluate whether DC-T cell interactions identified by LIPSTIC are cognate and dependent on MHC-II-antigen presentation by the DCs, we reconstituted lethally irradiated C57BL/6J mice with a mixture of *Cd40*^G5/G5^ and *Cd40*^G5/G5^,*H2*^−/−^ (MHC-II-deficient) bone marrow (**Fig. 1d**). Labeling was only observed in DCs that expressed MHC-II, confirming that the interactions captured by LIPSTIC are dependent on cognate antigen presentation (**Fig. 1e**). Functional *in vitro* differentiation co-cultures where 150 *Cd40*^G5/G5^ gLN DCs are incubated with 750 naïve CD4^+^ OT-II CFSE-labeled T cells, showed that biotin^+^, but not biotin^−^, DCs are able to induce T cell proliferation to a level above background as observed with gLN DCs from OVA-naïve mice (**Fig. 1f**). When we added exogenous OT-II peptide to the cultures, both biotin^+^ and biotin^−^ DCs induced T cell proliferation; however, only biotin^+^ DCs were capable of inducing Foxp3^+^ Treg differentiation (**Fig. 1g**). These data show that biotin^+^ DCs were loaded with OVA antigen *in vivo* and have an intrinsic ability to induce a regulatory T cell response. Corroborating previous studies^5,11^, LIPSTIC labeling was detected only on migratory (MHC-II^hi^) but not resident (MHC-II^int^) DCs (**Fig. 1h**). Despite higher frequency of total cDC1s compared to cDC2s in the proximal duodenal gLNs (D1 and D2; the preferential sites for food-specific pTreg induction^3^), labeling of cDC1 and cDC2 by OT-II T cells occurred at a similar ratio (**Fig. 1i**). Therefore, both major migratory subsets of DCs were equally capable of presenting dietary antigens to specific CD4^+^ T cells. We conclude that LIPSTIC is a sensitive and specific tool for monitoring intestinal cell-cell receptor-ligand interactions *in vivo* which accurately identifies the DCs that prime dietary antigen-specific CD4^+^ T cells in the gLNs.

To gain further insight into the profile of food antigen-presenting DCs, we performed plate-based single-cell mRNA sequencing (sc-RNA-seq) of biotin^+^ and biotin^−^ DCs from the duodenal gLNs (D-gLNs) 24 h after an i.g. administration of OVA. DCs fell into four transcriptional clusters (**Fig. 2a**), two corresponding to migratory DCs (Clusters 0 and 1) and two that corresponded to resident DCs (Clusters 2 and 3) based on expression of *Ccr7* and MHC-II (**Suppl. Fig. 2a, b**). Among migratory DCs, Cluster 0 expressed signatures associated with a cDC2 phenotype, while Cluster 1 DCs were associated with a cDC1 phenotype (**Suppl. Fig. 2c-e**). In agreement with flow cytometry data, biotin^+^ DCs comprised a mixture of migratory cDC1s and cDC2s at a roughly 1:1 ratio (**Fig. 2b, c**). Differential gene expression comparing biotin^+^ and biotin^−^ cDC1s and cDC2s from the D-gLNs showed that both subsets of biotin^+^ DCs upregulated genes encoding for markers of DC maturation (*Cd80, Cd86, Cd40*) and the cytokine *Ebi3* (**Fig. 2d**). Flow cytometry analysis confirmed higher expression of surface CD40, CD80, and CD86 in biotin^+^ DCs (**Suppl. Fig. 2f-h**). cDC1s showed increased expression of several genes associated with pTreg generation, such as *Aldh1a2* (RALDH2), *Tgfb2* (cytokine TGF-β2), *Itgb8* (integrin β8), *Ncoa7* and *Sod1^20,21^* (**Fig. 2e and Suppl. Fig. 2e**). In contrast, cDC2s were characterized by high expression of genes associated with inflammatory responses (*Cd44, Tyrobp, Bhlhe40, Fcer1g*)^22–25^ (**Fig. 2f**). To evaluate the ability of each biotin^+^ DC subset to induce pTregs, we co-cultured biotin^+^ cDC1s or biotin^+^ cDC2s with naïve CD4^+^ OT-II T cells. This showed that only biotin^+^ cDC1s were capable of inducing pTreg differentiation above background levels (**Fig. 2g**). Our data therefore suggest that DCs actively engaged in priming food-specific CD4^+^T cells display an activated profile and, even though cDC2 are able to present dietary antigens, pTreg differentiation is predominantly dependent on cDC1.

**Figure 2.**
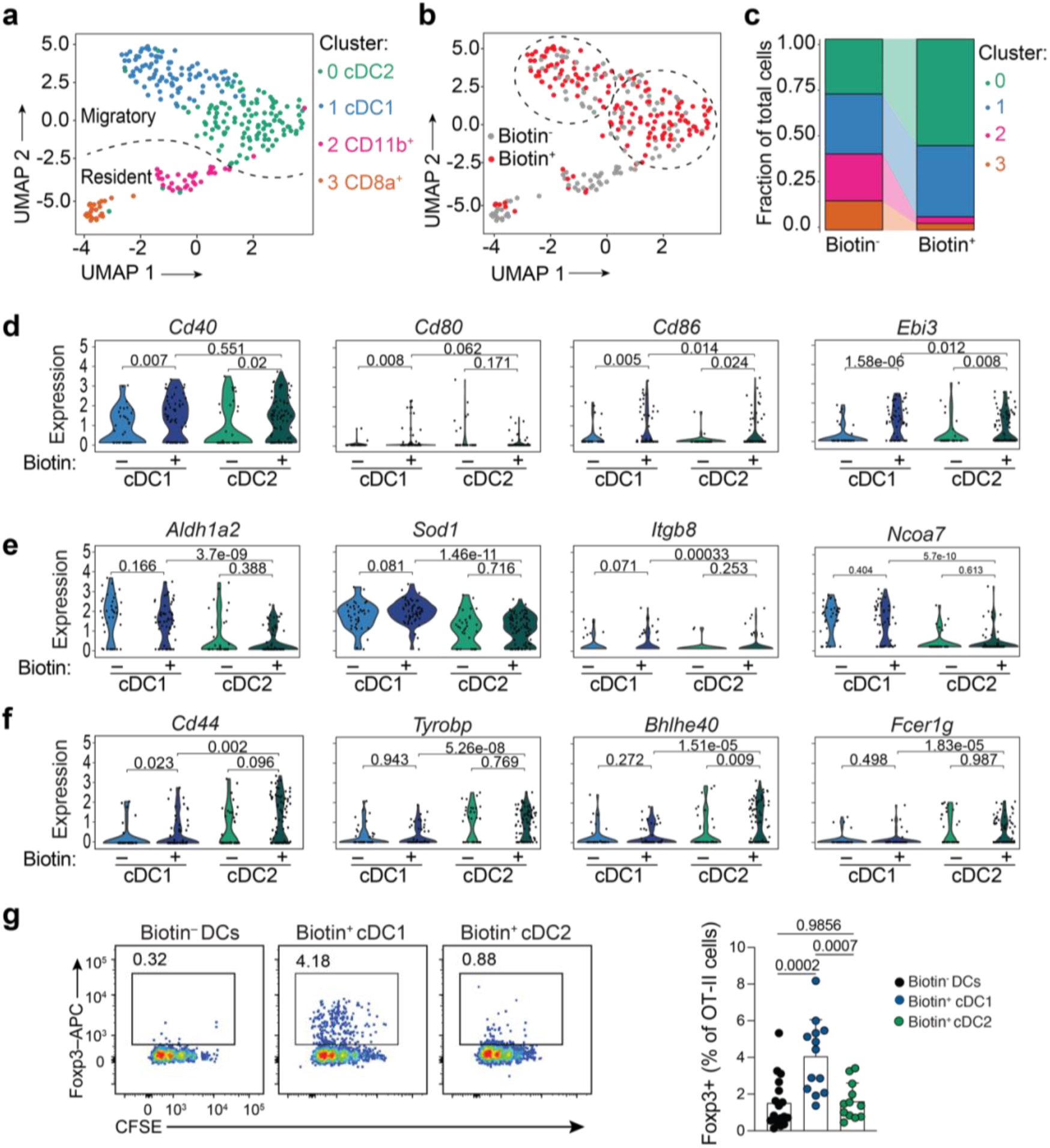
Biotin^+^ cDC1 promotes food-specific pTreg differentiation. **(a-f)** CD45.2 *Cd40*^G5/G5^ mice were adoptively transferred with 1 × 10^6^ naive CD45.1 CD4^+^*Cd40lg^SrA/Y^* OT-II T cells prior to one dose of intragastric OVA. Cell-cell interaction was revealed by LIPSTIC protocol 24 h later. Biotin^−^ and biotin^+^ D-gLNs DCs were single-cell sorted and subjected to sc-RNA-seq. **(a)** Uniform manifold approximation and projection (UMAP) plot. Cells were pooled from 7 mice from 2 independent experiments. Dotted line indicates the location of resident/migratory DC boundary. **(b)** Distribution of biotin^−^ (grey) and biotin^+^ (red) DCs. Dotted lines indicate the location of Cluster 0 and 1. **(c)** Proportion of cells in each transcriptional cluster among biotin^−^ and biotin^+^ DCs. **(d)** Expression of genes significantly upregulated in both biotin^+^ cDC1 and cDC2 compared to biotin^−^ cDC1 and cDC2. **(e, f)** Expression of genes significantly upregulated exclusively in **(e)** cDC1 or **(f)** cDC2. **(g)** OT-II CFSE-labeled T cells were co-cultured *in vitro* with sorted biotin^−^ DCs, biotin^+^ cDC1s or biotin^+^ cDC2s from D-gLNs in the presence of exogenous OT-II peptide for 96 h. Representative flow plots showing percentage of Foxp3^+^ cells among proliferated OT-II T cells (left), and quantification of data (right). Each dot represents one mouse (n = 3, 4 mice per group, pool of four independent experiments). *P*-values were calculated by one-way ANOVA and Wilcoxon signed-rank test was used for (D, E and F).

We previously showed that infection with *Strongyloides venezuelensis (S.v.)*, a helminth with duodenal tropism, prevents localized food-specific pTreg development and impairs induction of oral tolerance by generating an immunological conflict at the site of oral antigen presentation, the D-gLNs^3^. To determine whether any helminth infection can interfere with food-specific pTreg differentiation and oral tolerance, we infected C57BL/6 mice with *Heligmosomoides polygyrus (H.p.)*, which also displays duodenal tropism^26^. Five days post-infection (d.p.i), we adoptively transferred naïve OT-II T cells into host mice, followed by two doses of i.g. OVA (**Fig. 3a**). OT-II pTreg induction was only slightly impacted by *H.p*. infection compared to the drastic reduction in pTregs induced by *S.v*. (**Fig. 3a, b**). Accordingly, oral tolerance was intact following *H.p*.infection (**Suppl. Fig. 3a-d**), suggesting that the pTregs generated by OVA feeding were sufficient to prevent overt inflammation upon OVA-adjuvant immunization^5^. Both *H.p*. and *S.v*. infections led to decreased in total cDC1 and increased of total cDC2 frequencies in the D-gLNs, which was more extreme in *S.v*. infected mice (**Fig. 3c, d**).

**Figure 3.**
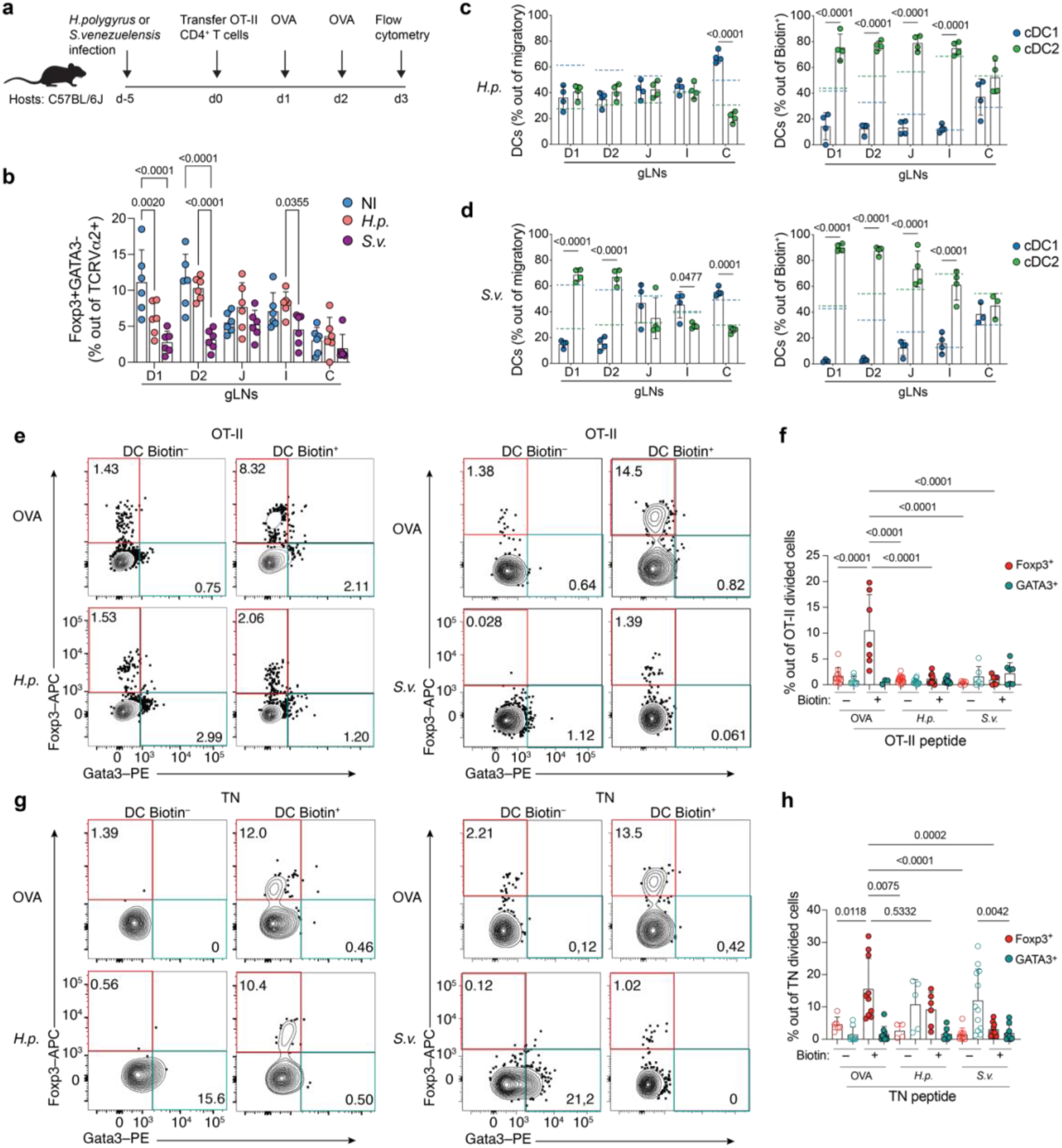
Helminthic infections affect food-specific pTreg induction by DCs. **(a, b)** CD45.2 C57BL/6 mice were infected with *S. venezuelensis (S.v.)* or *H. polygyrus (H.p.)* 5 days prior adoptively transfer of 1 × 10^6^ naive CD45.1 CD4^+^ OT-II T cells. Mice received two doses of intragastric OVA 48 h and 24 h prior analysis. Non-infected mice (NI) were used as control. **(a)** Experimental setup for panel (b). **(b)** Percentage of Foxp3^+^ cells among CD45.1 TCRVα2^+^ (OT-II) T cells in gLNs (n = 3 mice per group, pool of two independent experiments). **(c-h)** CD45.2 *Cd40*^G5/G5^ mice were adoptively transferred with 1 × 10^6^ naive CD45.1 CD4^+^*Cd40lg*^SrtA/Y^ OT-II T cells prior to one dose of intragastric OVA. Cell-cell interaction was revealed by LIPSTIC protocol 24 h later. Non-infected mice (OVA) were used as control. **(c, d)** Percentage of cDC1 and cDC2 out of total migratory DCs (left) and percentage of cDC1 and cDC2 out of biotin^+^ DCs (right) of **(c)** *H.p-* or **(d)** *S.v*-infected mice. Dashed lines show mean value of cDC1 (blue) or cDC2 (green) percentage at steady state (OVA) as in Figure 1i. **(e-h)** Sorted biotin^−^ or biotin^+^ DCs from D-gLNs of non-infected mice (OVA) or mice infected with *H.p* (left) or *S.v* (right) were co-culture with OT-II CFSE-labeled T cells **(e, f)** or TN CFSE-labeled T cells **(g, h)** *in vitro* for 96 h. **(e, g)** Representative flow plots showing percentage of Foxp3^+^ and GATA3^+^ cells among proliferated T cells co-cultured with biotin^−^ or biotin^+^ DCs from OVA, *H.p* (left) or *S.v* (right) infected mice and **(f, h)** quantification of data. OT-II or TN peptide were added to the corresponding co-cultures wells. Each dot represents one mouse (n = 3, 4 mice per group, pool of three independent experiments). *P*-values were calculated by two-way ANOVA.

To determine how these distinct helminth infections affected DC presentation of dietary antigen to food-specific CD4^+^ T cells, we infected *Cd40*^G5/G5^ mice with *H.p*. or *S.v*. and then adoptively transferred naïve *Cd40lg*^SrtA/Y^ OT-II T cells at 5 d.p.i, administered i.g. OVA, and performed LIPSTIC labeling as described above. Whereas approximately 15% of DCs presenting dietary antigen to CD4^+^ T cells during *H.p*. infection were of the cDC1 phenotype, this fraction fell to virtually zero in the D-gLNs of *S.v*. infected mice (**Fig. 3c, d and Suppl. Fig. 3f-h**). To test the ability of OVA-presenting biotin^+^ DCs to induce pTregs in the context of these helminth infections, we co-cultured biotin^+^ or biotin^−^ DCs from *H.p* or *S.v*. infected mice with naïve CD4^+^ OT-II T cells in the presence of exogenous OT-II peptide. As observed previously (see Fig. 1e), whereas biotin^+^ DCs from uninfected mice were capable of inducing pTreg differentiation *in vitro*, biotin^+^ DCs from infected mice failed to induce pTregs (**Fig. 3e, f**). To assess the intrinsic ability of OVA-presenting biotin^+^ DCs to induce pTregs in a non-OT-II system, independently of the presence of antigen acquired *in vivo*, we performed co-culture experiments using monoclonal transnuclear (TN) T cells bearing an unrelated T cell receptor (TCR) specific for a peptide of the β-hexosaminidase enzyme of commensals of the Bacteroidetes phylum^27^. We co-cultured biotin^+^ and biotin^−^ DCs, labeled by the OT-II T cells *in vivo*, with naïve CD4^+^ TN T cells in the presence of the TN cognate peptide. TN cells differentiated into pTregs when cultured with biotin^+^ DCs from uninfected or *H.p*.-infected mice, but not from *S.v*.-infected mice (**Fig. 3g, h**). Thus, biotin^+^ DCs (labeled due to their ability to present OVA *in vivo*) were able to induce pTreg differentiation when presenting an unrelated antigen, confirming that this is an intrinsic trait of the biotin^+^ DCs rather than a property of the antigen they acquired *in vivo*. By contrast, biotin^−^ DCs from both helminth infections induced TN cells to differentiate towards a GATA3^+^ Th2 phenotype (**Fig. 3g, h**), in accordance with the *in vivo* findings of endogenous Th2 response to helminth infections^3,25^ (**Suppl. Fig. 3e**). Of note, *in vitro* pTreg induction of TN cells by biotin^+^ DCs derived from *H.p*.-infected mice and Th2 induction by biotin^−^ DCs under both helminth infections were not observed in OT-II cells, which may reflect different properties and activation thresholds of monoclonal T cells bearing different TCRs. We conclude that different helminth infections have distinct impacts on DCs presenting dietary antigen and inducing pTreg differentiation; even during these extreme type-2 immune contexts, DCs presenting dietary antigens did not promote overt diet-specific Th2 differentiation.

To better understand the influence of helminth infections on the phenotype of dietary antigen-presenting DCs, we next performed sc-RNA-seq profiling of D-gLNs biotin^+^ and biotin^−^ DCs in the context of *H.p* and *S.v*. infection and compared them to our dataset of OVA-presenting DCs in steady state (see Fig. 2). Transcriptional UMAP profiling assigned DCs to 6 major unbiased clusters, which we defined based on their top differentially expressed genes (**Fig. 4a, b**). Clusters 0, 1, 2 and 3 corresponded to migratory DCs, whereas Clusters 4 and 5 corresponded to resident DCs (**Fig. 4a-c**). Among migratory DCs, Cluster 0 cells expressed genes associated with a cDC1 signature whereas Clusters 1, 2 and 3 were associated with a cDC2 phenotype (**Suppl. Fig. 4a, b**), in accordance with previous reports showing that cDC2s display a more heterogeneous transcriptional program^28,29^. Among biotin^−^ DCs, we observed an expansion of Cluster 3 cDC2s during *H.p*. infection, which was even more pronounced in *S.v*.-infected mice (**Fig. 4b**). DCs from this cluster were characterized by high expression of genes associated with a pro-inflammatory Th2 response to worm infections, such as *Pdcd1lg2* (which encodes PD-L2)^30–32^, *Stat5a^33^, Ccl24* (a chemotactic factor for eosinophils and lymphocytes)^34^, and *Cd1d1* (involved in lipid-based antigen presentation to iNKT cells, which play a role in the response to worm infections^35^).

**Figure 4.**
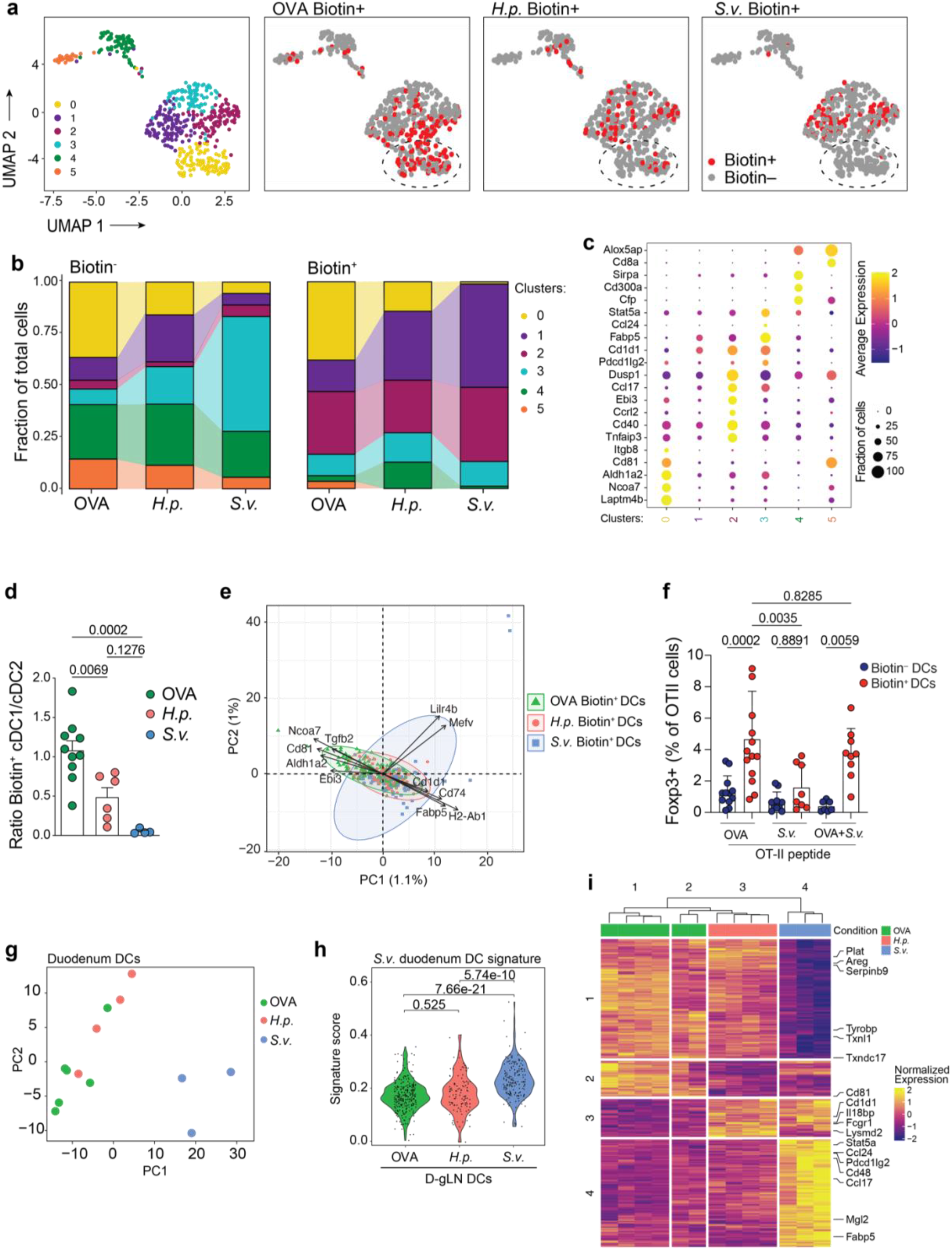
Helminth infection alter the profile of dietary antigen-presenting biotin^+^ DCs. CD45.2 *Cd40*^G5/G5^ mice were infected with *S. venezuelensis (S.v.)* or *H. polygyrus (H.p.)* 5 days prior adoptive transfer of 1 × 10^6^ naive CD45.1 CD4^+^*Cd40lg*^SrtA/Y^ OT-II T cells. Animals received 1 dose of intragastric OVA 18 h after OT-II T cells transfer. Cell-cell interaction was revealed by LIPSTIC protocol 24 h after OVA administration. Non-infected mice (OVA) were used as control. **(a)** UMAP plot showing clustering of DCs sorted from D-gLNs of non-infected mice (OVA group) or mice infected with *H.p* or *S.v*. Cells were pooled from 4-7 mice from 2 independent experiments. Distribution of biotin^−^ (grey) and biotin^+^ (red) DCs in the same plot. Dotted lines indicate the location of Cluster 0. **(b)** Proportion of cells in each transcriptional cluster among biotin^−^ and biotin^+^ DCs. **(c)** Dot plot showing expression of genes differentially expressed between Clusters. **(d)** Ratio of cDC1/cDC2 among biotin^+^ DCs from D-gLNs of non-infected mice (OVA) or mice infected with *H.p* or *S.v*. **(e)** Principal component analysis (PCA) of OVA, *H.p* and *S.v* biotin^+^DCs. **(f)** Percentage of Foxp3^+^ cells among proliferated OT-II CFSE-labeled T cells *in vitro* after 96 h of co-culture with D-gLNs OVA or *S.v* biotin^−^ or biotin^+^ DCs or a combination of OVA and *S.v* DCs. OT-II peptide was added to the co-cultures. Each dot represents one mouse (n = 3, 4 mice per group, pool of three independent experiments). **(g)** RNA-seq PCA of sorted DCs from duodenum lamina propria of C57BL/6 non-infected mice (OVA group) or mice infected with *H.p* or *S.v*. Cells were pooled from 4 and 7 mice per group from 2 independent experiments. **(h)** *“S.v*.duodenum DC” signature from (g) applied to single-cell D-gLN DC LIPSTIC transcriptomic dataset in (a). **(i)** Heatmap showing differentially expressed genes in OVA, *H.p* or *S.v* duodenum lamina propria DCs as in (g). *P*-values were calculated by one-way or two-way ANOVA and Wilcoxon signed-rank test.

Together with the induction of Th2 by biotin^−^ DCs (see Fig. 3g, h), these data suggest that Cluster 3 cDC2s were responsible for inducing a Th2 response against helminth infections, in agreement with previous studies showing that cDC2s are required for type-2 immunity to *H.p*. infection^25^. However, we did not observe any differences in the frequency of Cluster 3 cells among biotin^+^DCs in the three experimental groups, suggesting an absence of food-specific Th2 responses during helminth-induced type 2 immunity. Instead, there was a gradual increase in Cluster 1 cDC2s among DCs presenting dietary antigens in the context of *H.p* and especially *S.v*. infections (**Fig. 4b**). These DCs showed a less mature phenotype when compared to DCs from Cluster 0 or 3 (**Suppl. Fig. 4c, d)**. Cluster 1 DCs also lacked the tolerogenic potential of cDC1s (**Fig. 4c** and **Suppl. Fig. 4c)** but did not express the full pro-inflammatory signature of Cluster 3 DCs (**Fig. 4c** and **Suppl. Fig. 4d)**. This suggests that, during tolerance-disrupting infections, dietary antigen-presenting DCs fail to acquire Treg inducing capacity; yet, they do not promote overt Th2 differentiation towards the diet. This conclusion is supported by the *in vitro* experiments, where no Th2 induction was observed by DCs presenting dietary antigen (biotin^+^ DCs), even in the context of helminth infection (see Fig. 3e-h).

DCs from Cluster 0 displayed a tolerogenic transcriptional program (**Fig. 4c**) corresponding to that of the pTreg-inducing cDC1 population identified in Fig. 2, above. Importantly, the ratio of biotin^+^cDC1 to cDC2, approximately 1 at steady state (OVA), fell to 0.5 upon *H.p*. infection and to close to 0 in the presence of *S.v*. (**Fig. 4d,** see Suppl. Fig. 3g, h). In agreement, principal component analysis (PCA) of total biotin^+^ DCs under these three different conditions showed that *S.v*.infection markedly shifted the profile of DCs presenting dietary antigens away from the tolerogenic transcriptional program of cDC1s (**Fig. 4e**), with *H.p*. biotin^+^ DCs displaying an intermediate profile between OVA and *S.v*. biotin^+^ DCs (**Fig. 4b, e**). We therefore sought to understand whether the absence of dietary antigen presentation by biotin^+^ cDC1 (Cluster 0) or the expansion of biotin^+^ cDC2 (Cluster 1) presentation would be responsible for preventing the generation of pTreg in the context of *S.v*. infection. We co-cultured naïve CD4^+^ OT-II T cells with *S.v*. biotin^+^ DCs and OVA biotin^+^ DCs, either alone or in combination. Whereas *S.v*. biotin^+^ DCs failed to induce differentiation of pTregs, both OVA biotin^+^ DCs and a combination of *S.v*. and OVA DCs were able to do so (**Fig. 4f**). Thus, OVA biotin^+^ DCs are dominant in their ability to induce pTreg differentiation, indicating that the presence of a small fraction of cDC1s is both necessary and sufficient to induce a tolerogenic response even in the presence of a large pro-inflammatory DC population. This conclusion is supported by the finding that a reduced, but still detectable, dietary antigen presentation by cDC1 in the context of *H.p*. infection is sufficient to induce oral tolerance (see Fig. 4b, d and Suppl. Data Fig. 3a-c).

Lastly, we sought to determine whether the changes in DC populations and profiles observed in the D-gLNs of mice during helminth infections were imprinted already in the gut tissue, prior to DC migration to the gLNs. DCs from the duodenum lamina propria of uninfected, *H.p*.- or *S.v*.-infected mice were sorted at 7 d.p.i., 1 day after i.g. OVA administration. Transcriptional profiling of lamina propria DCs revealed that DCs from *S.v*.-infected mice clustered separately from those of steady-state (OVA) or *H.p*.-infected mice (**Fig. 4g**). We then selected significantly expressed genes exclusively detected in DCs from the duodenum lamina propria of *S.v*. infected mice to generate a *“S.v*. duodenum DC” signature, which we applied to our single-cell D-gLNs DC LIPSTIC transcriptomic dataset. D-gLNs DCs from *S.v*.-infected mice were enriched for the *S.v*.duodenum DC signature, suggesting that changes in DCs program were imprinted in the intestine, prior to their arrival at the gLNs (**Fig. 4h, i**). Furthermore, the pro-inflammatory Th2 inducing cDC2 signature observed in Cluster 3 of D-gLNs was enriched in the duodenum DCs of *S.v*.compared to OVA mice (**Suppl. Fig. 4e, f)**. Together, these data suggest that *S.v*. infection affects cDC differentiation, favoring a cDC2 in detriment of a cDC1 program in the lamina propria. This shift results in increased cDC2 migration and impaired pTreg generation in the D-gLNs and ultimately in failure to induce oral tolerance.

## Discussion

The ability to isolate the exact DCs within a population that present antigens to the T cells of interest *in vivo* has been a longstanding challenge to the field. Intestine-adapted LIPSTIC allowed us to identify individual DCs actively engaged in presenting dietary antigen and priming of food-specific CD4^+^ T cells in the gLNs. Hence, we were able to define the transcriptional program and function of these DCs at steady state and in the context of infection-mediated disruption of pTreg and oral tolerance induction.

Our study revealed that, although both cDC1s and cDC2s present dietary antigens, pTreg differentiation is predominantly dependent on cDC1s. These findings add to the concept of hierarchical roles for cDC subsets in pTreg induction, shown previously using mouse models of lineage ablation^5^. Whereas genetic models are limited by incomplete depletion of cDC1s as well as off-target effects, the maintenance of tolerance upon ablation of cDC1s by excising IRF8 in cDC precursors observed in our previous study^5^ resembles the effects of *H.p*. infection observed here: decreased, but not absent, cDC1s and dietary antigen-specific pTreg induction, but sustained oral tolerance. In contrast, *S.v*. infection prevents oral tolerance induction by completely abrogating cDC1 presentation of dietary antigens and pTreg generation. It remains to be determined whether additional “immunological conflicts” such as reovirus infection, shown to induce food-specific Th1 responses^36,37^, or scarring post bacterial infection, shown to prevent Treg and oral tolerance induction^38^, specifically interfere with dietary antigen presentation by cDC subsets. LIPSTIC allowed us to uncover a compartmentalized antigen presentation dynamics that prevents the induction of food-specific Th2 cells during robust type-2 immunity induced by helminth infections. Furthermore, the intestine-adapted LIPSTIC technology presented here might be useful for future investigations of *in vivo* DCs inducing regulatory and inflammatory T cell responses to commensals or enteric pathogen-derived antigens. In conclusion, this work provides insight into the role of cDC1s as the primary population responsible for inducing food-specific pTregs and oral tolerance, and uncovers a novel mechanism by which infection can disrupt this pathway to impair oral tolerance by exclusion of migratory cDC1s from the draining LNs, rather than by their reprogramming^36,39^. Our findings demonstrate how dynamic DC-T cell interactions drive tolerance or inflammation towards dietary proteins in the complex gut environment.

## Methods

### Mice

CD45.2 (C57BL6/J) and H2-/- mice were purchased from the Jackson Laboratories (strain numbers 000664 and 003584) and CD45.1 OT-II TCR-transgenic were originally purchased from Taconic Farms, Rag1-/-was bred out, and mice were maintained in our facilities (strain number 4234-M). *Cd40*^G5/G5^ and *Cd40lg*^SrtA/Y^ mice were generated and maintained in our laboratory^18^. Transnuclear (TN) mice were generated as described^40^ and maintained in our facilities. Male and female mice at 7-12 weeks old were used throughout the study. Mice were maintained at the Rockefeller University animal facilities under specific pathogen-free conditions. All protocols were approved by the Rockefeller University Institutional Animal Care and Use Committee.

### Reagents

Ovalbumin (grade III, A5378; grade VI, A2512) and Pyrantel Pamoate (P6210) were purchased from Sigma. LPS-free ovalbumin was from Hyglos, Germany (Cat. no 77161).

### Antibodies, staining and flow cytometry

Fluorescent-dye-conjugated antibodies were purchased from BD (USA) (anti-CD45.2, 560693; anti-CD103, 557495; anti-SiglecF, 552126; anti-CD40, 562846; anti-I-A/I-E (MHCII), 743876; anti-CD11c, 741139; anti-CD8a, 752640), eBioscience (USA) (anti-B220, 48-0452-82; anti-CD4, 83-0042-42; anti-CD8a, 56-0081-82; anti-CD11b, 47-0112-82; anti-CD11c, 25-0114-82, 17-0114-82 and 56-0114-82; anti-CD45.1, 25-0453-82; anti-FOXP3, 17-5773-82; anti-GATA3, 12-9966-42; anti-I-A/I-E (MHCII), 46-5321-82 and 56-5321-82; anti-Ly6G, 48-5931-82; anti-Va2, 48-5812-82), or Biolegend (USA) (anti-CD8a, 100744; anti-CD11b, 101236; anti-CD64, 139306; anti-TCRβ, 109220; anti-CD86, 105037; anti-CD80, 104722; anti-CD40L, 106510). Anti-biotin-PE antibody was purchased from Miltenyi Biotec (130-113-291) for biotin–LPETG SrtA substrate staining. Biotinylated anti-CD40L antibody was purchased by eBioscience (13-1541-82). Biotinylated anti-IgG1, Bethyl, A90-105B was purchased from R&D Systems, BAF566. Unconjugated anti-OVA IgG1, was purchased by Biolegend, 520501. Horseradish peroxidase conjugated Streptavidin was purchased from Jackson Immuno Research Laboratories, Inc. Aqua LIVE/DEAD® Fixable Aqua Dead Cell Stain Kit, L-34965, and Cell Trace CFSE Proliferation kit (C34554) was purchased from Life Technologies. Zombie NIR™ Fixable Viability kit, 423106, was purchased from Biolegend. Cell populations were stained with Aqua or Zombie in PBS, followed by incubation with Fc block and antibodies against the indicated cell surface markers in FACS buffer (PBS, 1% BSA, 10 mM EDTA, 0.02% sodium azide). The cells were analyzed live or fixed in 1% PFA/PBS. For intracellular staining, cells were first stained for surface epitopes and then fixed, permeabilized and stained according to the manufacturer’s protocol (eBioscience 00-5123-43). Flow cytometry was performed on FACSymphony A5 (BD Biosciences) and analyzed using FlowJo Software package (Tri-Star).

### SrtA substrate

Biotin–aminohexanoic acid-LPETGS (C-terminal amide, 95% purity) was purchased from LifeTein (custom synthesis) and stock solutions prepared in PBS at 20 mM.

### LIPSTIC *in vivo*-labeling experiments

Biotin–LPETG substrate was injected into *Cd40*^G5/G5^ mice intraperitoneally (i.p.) (100 μl of 20 mM solution in PBS) six times 20 min apart. gLNs were collected 40 min after the last injection. Mice were briefly anaesthetized with isoflurane at each injection. For CD40L-blockade experiments, mice were injected intravenously with 200 μg of CD40L-blocking antibody (clone MR-1, BioXCell) 22 h prior to substrate injection.

### Lymphocyte and dendritic cell isolation from lymph nodes

Tissues were dissected into cold RPMI supplemented with 10% heat-inactivated fetal bovine serum (Hyclone), 2 mM L-glutamine, 100 units per ml of penicillin, 100 μg/ml of streptomycin sulfate, 1 mM sodium pyruvate, 0.1 mM non-essential amino acids, 10 mM HEPES (all from Gibco), and 50 μM β mercaptoethanol (Sigma). Lymph nodes were finely chopped and incubated in 400 U/ml Collagenase D (Roche) in supplemented RPMI for 25 min at 37°C, 5% CO2. Single cell suspensions were extracted from connective tissue by taking up and resuspending the digests five times.

### Segmentation of gut-draining lymph nodes

The mouse gLNs consist of one hepatic–coeliac lymph node co-draining the duodenum, pancreatic–duodenal lymph nodes draining the duodenum and separately the ascending and transverse colon, the main mLN chain draining the distal duodenum, jejunum, ileum, caecum and proximal ascending colon, and the caudal and iliac lymph nodes draining the descending-distal colon. Mesenteric lymph nodes draining intestinal segments were identified anatomically by following the lymphatic vessels connecting the colon, ileum and jejunum to their lymph nodes. Duodenal lymph nodes were revealed by gavaging with 100 μl of olive oil (Sigma) and determining the most stomach-proximal lymph nodes surrounded by chyle, indicative of duodenal drainage, 1 h after gavage, as described before^3^.

### Dendritic cells isolation from duodenum lamina propria

Intestines were separated from mesentery, and Peyer’s patches and faeces were removed. For segmentation of the small intestine, the upper 25% of the small intestine was taken as duodenum as described before^3^. Intestines were cut longitudinally and washed twice in PBS. Tissue was cut into 1-cm pieces, mucus was removed by incubating the tissue for 10 min in PBS and 1 μM DTT, and the epithelium was removed by two incubations in 25 ml of RPMI, 2% FCS and 30 mM EDTA for 10 min at 37 °C at 230 r.p.m. with vigorous shaking after each incubation. Tissues were washed in PBS over a sieve, then finely chopped and digested in 6 ml of RPMI, 2% FCS, 200 μg/ml DNaseI (Roche) and 2 mg/ml collagenase 8 (Gibco) for 45 min at 37 °C, 5% CO2. Digests were taken up and resuspended 10 times and passed through a sieve, and the collagenase was quenched by addition of 15 ml cold RPMI, 2% FCS. Cell pellets were resuspended in 40% Percoll (BD Pharmigen) complemented with RPMI, 2% FCS, passed through a 100-μm mesh and separated by centrifugation in a discontinuous Percoll gradient (80%/40%) at 1000g for 25 min at room temperature. DCs were isolated from the interphase, washed, and stained for FACS bulk sorting.

### Single-cell sorting

Dendritic cells were collected, as described above, from the duodenum-gLN of *Cd40*^G5/G5^ male mice (8-12 weeks old). Single cells from three to four mice per each condition were index-sorted directly into 96-well plates containing 5μl of TCL buffer (Qiagen) supplemented with 1%β-mercaptoethanol using a BD FACS Aria II instrument. After sorting, plates were immediately frozen on dry ice and stored at −80°C before processing. All sorting data were analyzed using FlowJo software v.10. Dendritic cells were sorted as Aqua^−^CD45.2^+^CD45.1^−^Lin^−^ (TCRβB220^−^CD64^−^)CD11c^hi^MHCIΓ^nt/hi^Biotin^−^ or Biotin^+^. Dendritic cells were also stained for, but not sorted based on, CD103, CD11b and CD8α for index analysis.

### Bulk sorting

Dendritic cells were collected, as described above, from the duodenum lamina propria of C57BL/6 male mice (8-12 weeks old). Three hundred cells from three to six mouse per condition were sorted directly into 25 μl TCL buffer (Qiagen, 1031576) supplemented with 1%β-mercaptoethanol at single cell precision using a BD FACS Aria II instrument. After sorting, samples were immediately frozen on dry ice and stored at −80°C before processing. Dendritic cells were sorted as Aqua^−^CD45.2^+^Lin^−^(TCRβ^−^B220^−^CD64^−^)CD11c^hi^MHCII^int/hi^.

### Library preparation for single cell RNAseq

Each sorted plate contained all conditions assayed in each replicate and libraries were prepared as previously described^41^. RNA was extracted from single cells using RNAClean XP Solid Phase Reversible Immobilization (SPRI) beads (Agentcourt, Beckman Coulter), and hybridized first using RT primer (5′/5Biosg/AAGCAGTGGTATCAACGCAGAGTACTTTTTTTTTTTTTTTTTTTTTTTTTTTTTTVN-3′) and then reverse transcribed into cDNA using TSO primer (5′-AAGCAGTGGTATCAACGCAGAGTACATrGrGrG-3’) and RT maxima reverse transcription (Thermo Fisher Scientific). cDNA was amplified using ISPCR primer (5′-AAGCAGTGGTATCAACGCAGAGT-3′) and KAPA HiFi HotStart ReadyMix (Thermo Fisher Scientific), cleaned up using RNAClean XP SPRI beads three times, and tagmented using Nextera XT DNA Library Preparation Kit (Illumina) following manufacturer’s instructions. For each sequencing batch, up to four plates were barcoded at a time with Nextera XT Index Kit v2 Sets A–D (Illumina). Finally, barcoded libraries were pooled and sequenced using Illumina Nextseq 550 platform.

### Library preparation for bulk RNA-seq

RNA was isolated using RNAClean XP beads (Agentcourt, A63987) on a magnetic stand (DynaMag, Invitrogen 12331D). Reverse transcription primers were: P1-RNA-TSO: Biot-rArArUrGrArUrArCrGrGrCrGrArCrCrArCrCrGrArUrNrNrNrNrNrNrGrGrG; P1-T31: Biot-AATGATACGGCGACCACCGATCG31T; P1-PCR: Biot-GAATGATACGGCGACCACCGAT. RNA was eluted for 1 min at room temperature in cDNA synthesis mix 1 (0. 5 μl P1-T31 (20 μM), 0.3 μl RNasin plus (Promega, N2615), 1.5 μl 10 mM dNTP, 3.5 μl 10 mM Tris pH 7.5, 0.5% IGEPAL CA-630 (Sigma) and 1.7 μl RNase free ddH2O) and the beads pipetted up and down ten times. The eluted sample was then incubated for 3 min at 72 °C, followed by 1 min on ice, then 7.5 μl of mix 2 was added (3 μl 5Å~ FS Buffer SS, 0.375 μl 100 mM DTT, 0.375 μl RNasin plus, 0.5 μl P1-RNA-TSO (40 μM), 0.75 μl Maxima RT Minus H (Thermo Scientific, EP0751), 1.8 μl 5M Betaine (Sigma, B0300), 0.9 μl 50 mM MgCl2 and 0.175 μl RNase free ddH2O. Reverse transcription occurred during a thermal cycle of one cycle (90 min at 42 °C), 10 cycles (2 min at 50 °C, 2 min at 42 °C) and one cycle (15 min at 70 °C), and the product was kept at 4 °C. The cDNA was then amplified using 15 μl of the reverse transcription product, 20 μl 2Å~ KAPA HiFi HS Ready Mix (Kapabiosystems, KK2601), 1.5 μl P1-PCR (10 μM), and 3.5 μl RNase-free ddH2O. Amplification occurred during the following cycles: one cycle of 3 min at 98 °C, 20 cycles of (15 s at 98 °C, 20 s at 67 °C, 6 min at 72 °C), and one cycle of 5 min at 72 °C, and the product was kept at 4 °C. PCR product (20 μl) was cleaned up using 16 μl RNAClean XP beads. The cDNA was eluted in 20 μl RNase-free ddH2O and kept at −20 °C. Isolated amplicons were confirmed to be 1,500–2,000 bp long by a High Sensitivity DNA Assay (Bioanalyzer). Concentration of all sample was measured on a Qubit fluorometer (Thermo Fisher), all samples were adjusted to 0.2 ng/μl with ddH2O, and 2.5 μl cDNA was subjected to Nextera XT DNA Library preparation (Illumina) using a Nextera XT Index Kit (Illumina, FC-131-1002) according to the manufacturer’s protocol, except that all volumes were used at 0.5× the indicated volumes. Sample quality was again verified by Bioanalyzer, and sample concentrations were measured on the Qubit fluorometer and adjusted to a concentration of 4.54 ng/μl. All samples were pooled at equal contributions and run in multiple lanes. Sequencing was performed using 75-base single-end reading on a NextSeq instrument (Illumina).

### Single-cell RNA-Seq analysis

Fastq sequence files generated from Smartseq2 libraries were aligned to the mouse genome (mm39) associated with the mouse transcriptome annotations (v. gencode M29) using STAR (v. 2.7.10a; Dobin et al., 2013). Subsequently, genome-mapped BAM files were processed through RSEM (v. 1.3.1; Li and Dewey, 2011) for gene quantification. The matrix of gene counts was then used as input for analysis by the R package Seurat (v. 4.1.2.; Stuart et al., 2019). Next, the dataset was normalized by the LogNormalize method implemented by Seurat. Finally, we “regressed out” the mitochondrial gene abundance and library size with the ScaleData function to control unwanted experimental noise sources. Additionally, cells containing more than 5% sequence reads aligned to mitochondrial genes were excluded before normalization. Single cells were clustered, and gene expression was evaluated with the Seurat workflow. Gene signature scoring was performed by using Seurat’s AddModuleScore. Heatmaps were produced with the ComplexHeatmaps package for R (10.1093/bioinformatics/btw313). Differentially expressed gene analyses between clusters or biological groups were performed by the Seurat function FindAllMarkers with the non-parametric Wilcoxon Rank sum test, and pvalues were ajusted using the Bonferroni correction. Genes were considered for downstream analysis when having a log2 fold change greater or smaller than 0.5 and exhibiting an adjusted p-value of at least 0.05.

### Bulk RNA-Seq Analysis

Raw fastq files were aligned and quantified with STAR (v. 2.7.10a) by using the mouse genome (mm39) and the mouse transcriptome (gencode M29) (v. 2.7.10a; Dobin et al., 2013). Next, the count matrix was imported to the R environment and processed by the DESeq2 package (v. 1.34) pipeline (doi:10.1186/s13059-014-0550-8). Briefly, genes from samples expressing less than ten reads were pre-filtered. Gene expression among samples was normalized by applying a negative binomial distribution model. The Wald test was employed to determine differential gene expression between conditions. Genes containing adjusted pvalues (FDR) less than 0.1 were considered for downstream analysis, and log2 fold changes were shrunk using the apeglm algorithm.

### Adoptive T cell transfer

Naive CD4^+^ T cells from spleen and lymph nodes were isolated by negative selection using biotinylated antibodies against CD8α, CD25, CD11c, CD11b, TER-119, NK1.1, and B220 and anti-biotin MACS beads (Miltenyi Biotec). The purity of transgenic CD4^+^OT-II T cells was verified by flow cytometry (CD45.1^+^Vα2^+^Vβ5^+^CD25^−^, typically >90%). 1 × 10^6^ OT-II cells were transferred by retro-orbital injection under isoflurane gas anesthesia.

### Oral antigen administration

OVA (grade III, Sigma, A5378) was administered intragastric at 50 mg in 200 μl PBS using plastic gavage needles. OVA was given 16-18 h after adoptive OT-II cell transfer.

### Bone marrow chimaeras

C57BL/6 recipient mice were lethally irradiated with two doses of 450 Rads given 4 h apart. After irradiation, recipients were reconstituted by intravenous injection of hematopoietic cells collected from femurs and tibiae of donor mice, as described before^18^. Mice were used for experiments 8–12 weeks after irradiation.

### *In vitro* dendritic cell-T cell co-culture

LIPSTIC substrate was delivered *in vivo* and dendritic cells were collected from the duodenum-gLN, as described above. For experiments on Fig. 1f, 150 Biotin’, Biotin^+^ or total DCs were sorted into U bottom 96well plate containing supplemented RPMI and 2% of T-stim media (VWR). 750 naïve CD4^+^ OT-II CFSE-labeled T cells from the spleen were sorted into the corresponding wells. Cells were incubated at 37°C, 5% CO2 for 96 h. To determine the number of proliferated cells, all cells in each well were recorded by FACS and gated based on CFSE staining. For all other co-culture experiments, 1 μM of OT-II peptide (chicken OVA amino acids 323-339 - Anaspec AS-27024) or TN peptide (β-hex peptide sequence YKGSRVWLN - GenScript)^27^ was added to the wells. 50 Biotin^−^ or Biotin^+^ DCs and 750 naïve CD4^+^ OT-II or TN CFSE-labeled T cells from the spleen were sorted into U bottom 96well plate containing supplemented RPMI and 2% of T-stim media. Cells were incubated at 37°C, 5% CO2 for 96 h, followed by surface and intracellular FACS staining, as described above. Freshly isolated congenic CD45.2^+^ splenocytes were added to the wells prior staining to prevent cell loss. Naïve CD4^+^ OT-II or TN T cells from the spleen were first isolated by negative selection and CFSE-labeled, as described above. These cells were then sorted as Aqua^−^CD45.1^+^TCRVα2^+^CD62L^+^CD44^−^CD25^−^. Dendritic cells were sorted as Aqua^−^CD45.2^+^CD45.1^−^Lin^−^(TCRβ^−^B220^−^CD64^−^)CD11c^hi^MHCII^int/hi^Biotin^−^ or Biotin^+^. cDC1s were sorted as Aqua^−^CD45.2^+^CD45.1^−^Lin^−^(TCRβ^−^B220^−^CD64^−^)CD11c^hi^MHCII^hi^CD103^+^CD11b^−^ and cDC2s were sorted as Aqua^−^CD45.2^+^CD45.1^−^Lin^−^(TCRβ^−^B220^−^CD64^−^)CD11c^hi^MHCII^hi^CD103^+/−^CD11b^+^.

### *Strongyloides venezuelensis* passage and infection

*S. venezuelensis* was maintained in our facility in NSG mice by subcutaneous infection with 1000 stage 3 (L3) larvae, resulting in chronic infection of this strain. For each experiment, feces of infected NSG mice were collected and spread on Whatman paper, which was placed into a beaker with water and incubated at 28C for 3 days. Mice were infected subcutaneously with 700 L3 larvae in 200 ml water per mouse. *S. venezuelensis* was passaged periodically by infecting naïve adult NSG mice.

### *Heligmosomoides polygyrus* infection

*H. polygyrus* larvae was kindly provided by William Gause (Rutgers University). Mice were infected by oral gavage with 125 third-stage larvae of *H. polygyurs* in 100 μl of PBS. For the oral tolerance experiments on Supplementary Figure 3, C57BL/6 mice were treated with two intragastric doses of Pyrantel Pamoate (300 μl of 10mg/ml in PBS) to clear *H. polygyurs* infection.

### Alum immunization and airway challenge

Twelve days after oral administration of OVA, 4 μg of endotoxin-free OVA antigen adsorbed to 40 μl Imject Alum Adjuvant (Fisher Scientific) was injected intraperitoneally in a final volume of 400 μl made up with PBS. Immunization was repeated after 7 days. To induce airway inflammation, mice were anaesthetized and intranasally administered 10 μg of sterile OVA grade VI in 50 μl PBS (25 μl per nostril) on days 14, 17 and 21 after the first intraperitoneal immunization.

### Bronchoalveolar lavage (BAL) and infiltrate analysis by flow cytometry

Mice were anaesthetized by intraperitoneal injection of 0.35 ml 2.5%avertin (Sigma), the trachea was cannulated and lungs were lavaged once with 0.5 ml and then with 1.0 ml PBS. Total BAL cells were counted after erythrocyte lysis and stained for FACS analysis. Lungs were perfused via the right ventricle with 10 ml saline to wash out residual blood. One lobe was digested in 400 U/ml collagenase D/RPMI and processed for FACS analysis. Eosinophils were determined as CD45^+^SSA^hi^MHCII^−^CD11b^+^Ly6G^int^SiglecF^+^.

### Anti-OVA IgG1 ELISA

Enzyme-linked immunosorbent assays (ELISAs) were performed as described previously^42^.

### Statistical analysis

Statistical analyses were performed in GraphPad Prism 9.0 software. Error bars indicate S.E.M. Comparisons between two treatment conditions were analyzed using two-tailed unpaired Student’s t-test. Wilcoxon signed-rank test was used for RNA-seq data (indicated in figure legends). Multivariate data were analyzed by applying one-way ANOVA or two-way ANOVA and Tukey’s multiple comparison post hoc test. A *P*-value of less than 0.05 was considered significant

## Data availability

The mouse sequencing data are available through the Gene Expression Omnibus under accession GSExxxx.

## Code availability

All code used for analysis in this manuscript is available at https://github.com/victoraLab/Lipstic_DC_Gut

## Acknowledgments

We thank all Mucida and Victora Lab members and Rockefeller University employees for their continuous assistance, particularly: A. Rogoz and S. Gonzalez for the maintenance of mice, RU Genomics core for assistance with sequencing, K. Gordon and K. Chhosphel for sorting, W. Gause for providing *H. polygyrus* larvae, M. London for critical reading of the manuscript. We also thank the Lafaille lab for fruitful discussions. This work was supported by a Pew Latin American postdoctoral fellowship (M.C.C.C.), a Life Sciences Research Foundation postdoctoral fellowship (T.B.R.C), The Howard Hughes Medical Institute, R01DK093674, R01DK113375, R21AI144827, and Food Allergy FARE/FASI Consortium (D.M.), DP1AI144248 (Pioneer Award), Burroughs-Wellcome PATH, Pew-Stewart Scholar, MacArthur Fellow (G.D.V.).

## Author contributions

M.C.C.C. initiated, designed, performed, and analyzed experiments and wrote the manuscript. A.L. and B.R. assisted with experiments. D.E. assisted with oral tolerance experiment in *H. polygyrus* infected mice. T.B.R.C performed RNAseq analysis. S.N.-H. generated the *Cd40lg*^SrtA^ mice. D.M. and G.D.V. conceived, initiated, designed, and supervised the research and wrote the manuscript. All authors revised and edited the manuscript and figures.

## Competing interests

G.D.V. has a U.S. patent on LIPSTIC technology (US10053683). G.D.V. is a scientific advisor for Vaccine Company Inc.

Correspondence and requests for materials should be addressed to M.C.C.C, G.D.V or D.M.

---

**Supplementary Figure 1.**
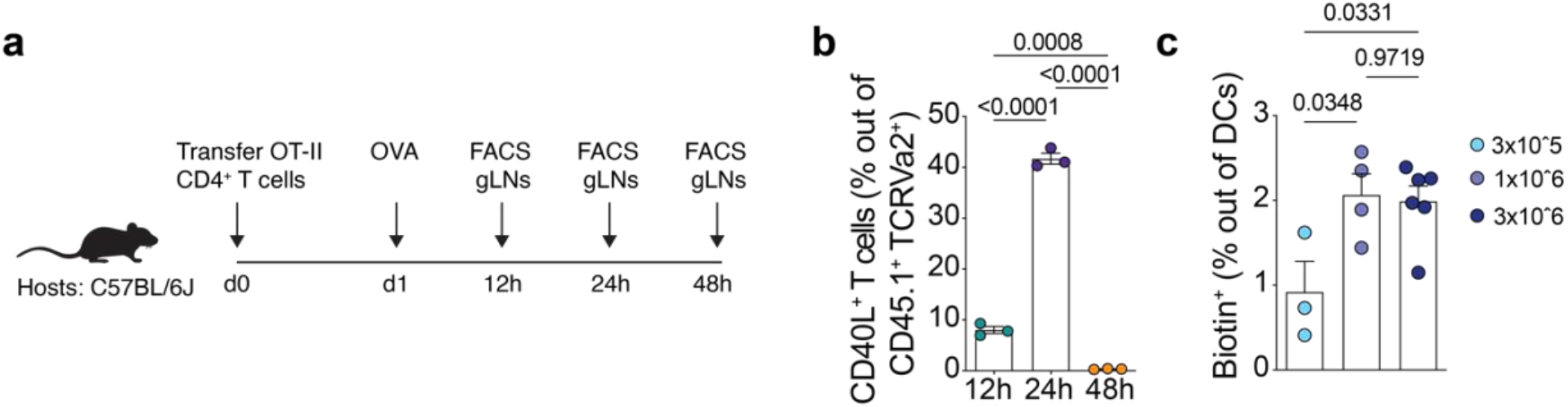
Intestine-adapted LIPSTIC optimization. **(a, b)** CD45.2 C57BL/6 mice were adoptively transferred with 1 × 10^6^ naive CD45.1 CD4^+^ OT-II T cells prior to one dose of intragastric OVA. Analyses were carried out at indicated time points. **(a)** Experimental setup for panel (b). **(b)** Percentage of CD40L^+^ T cells among CD45.1^+^TCRVα2^+^ cells in D-gLNs (n = 3 mice per group). **(c)** CD45.2 *Cd40*^G5/G5^ mice were adoptively transferred with 3 × 10^5^, 1 × 10^6^ or 3 × 10^6^ of naive CD45.1 CD4^+^*Cd40lg*^SrtA/Y^ OT-II T cells prior to one dose of intragastric OVA. Cell-cell interaction was revealed by LIPSTIC protocol 24 h later. **(c)** Percentage of labeled DCs in the D-gLNs (n = 3 to 6 mice per group). *P*-values were calculated by one-way ANOVA.

**Supplementary Figure 2.**
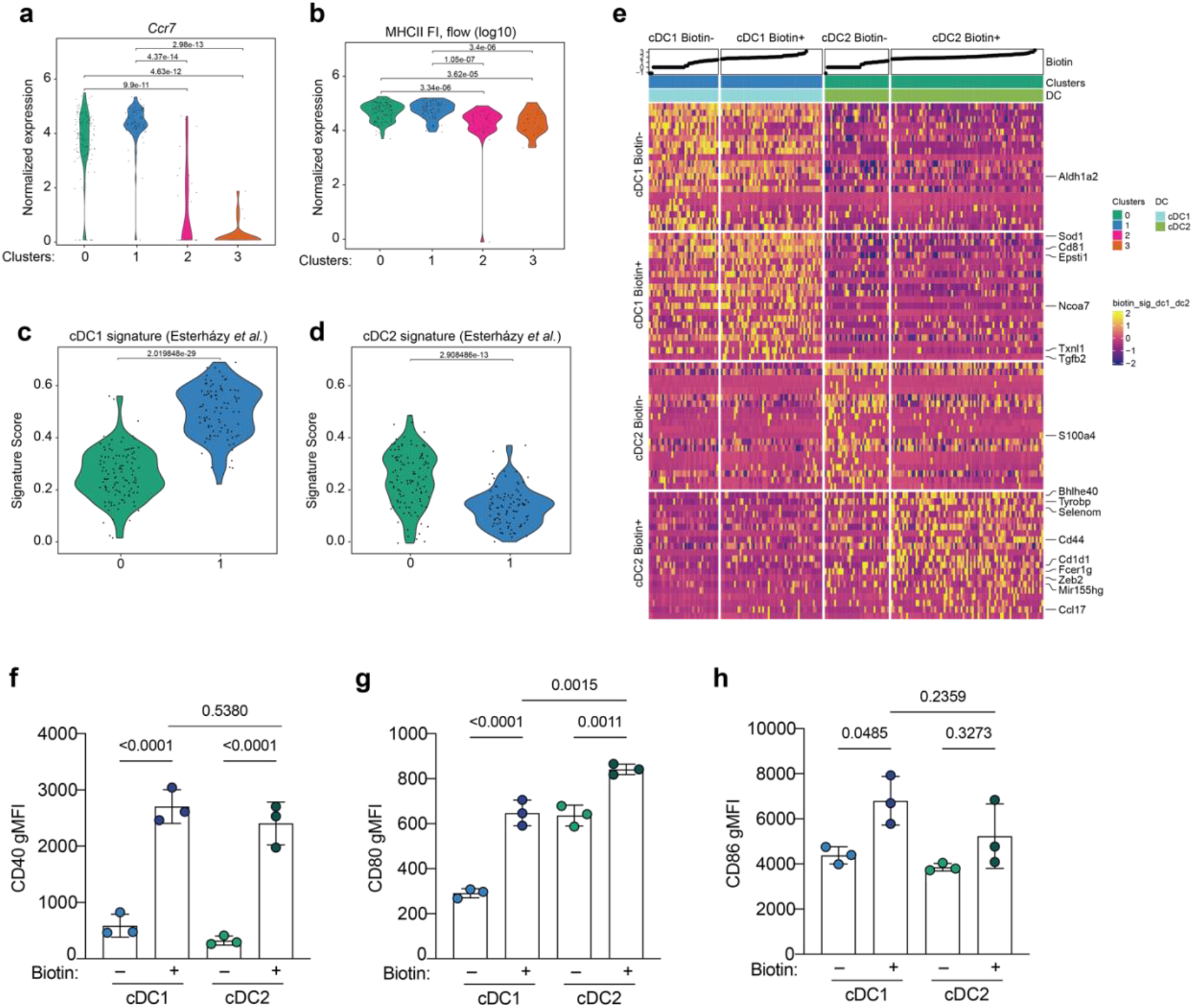
Single-cell transcriptomics of D-gLNs DCs. CD45.2 *Cd40*^G5/G5^ mice were adoptively transferred with 1 × 10^6^ naive CD45.1 CD4^+^*Cd40lg*^SrtA/Y^ OT-II T cells prior to one dose of intragastric OVA. Cell-cell interaction was revealed by LIPSTIC protocol 24 h later. D-gLNs biotin^−^ and biotin^+^ DCs were single-cell sorted and subjected to sc-RNA-seq. **(a)** *Ccr7* expression in different transcriptional Clusters as defined in Fig 2a. **(b)** Fluorescence intensity (FI) of MHCII staining in transcriptional Clusters as defined in Fig 2A. FI data was obtained from flow cytometry index-sorting files. Expression of **(c)** cDC1 and **(d)** cDC2 gene expression signatures, obtained from the literature^3^, in transcriptional Clusters 0 and 1. **(e)** Heatmap showing expression of genes significantly modulated in biotin^−^ and biotin^+^ cDC1 and cDC2. The uppermost rows show biotin FI from flow cytometry, followed by transcriptional clusters as shown in Fig. 2A and cDC1 or cDC2 as defined in (c and d). Selected genes are indicated in the right. **(f)** CD40, **(g)** CD80 and **(h)** CD86 geometric mean fluorescence intensity (gMFI) of biotin^−^ and biotin^+^cDC1s and cDC2s from D-gLNs (n = 3 mice per group). *P*-values were calculated by one-way ANOVA or Wilcoxon signed-rank test.

**Figure 1:**
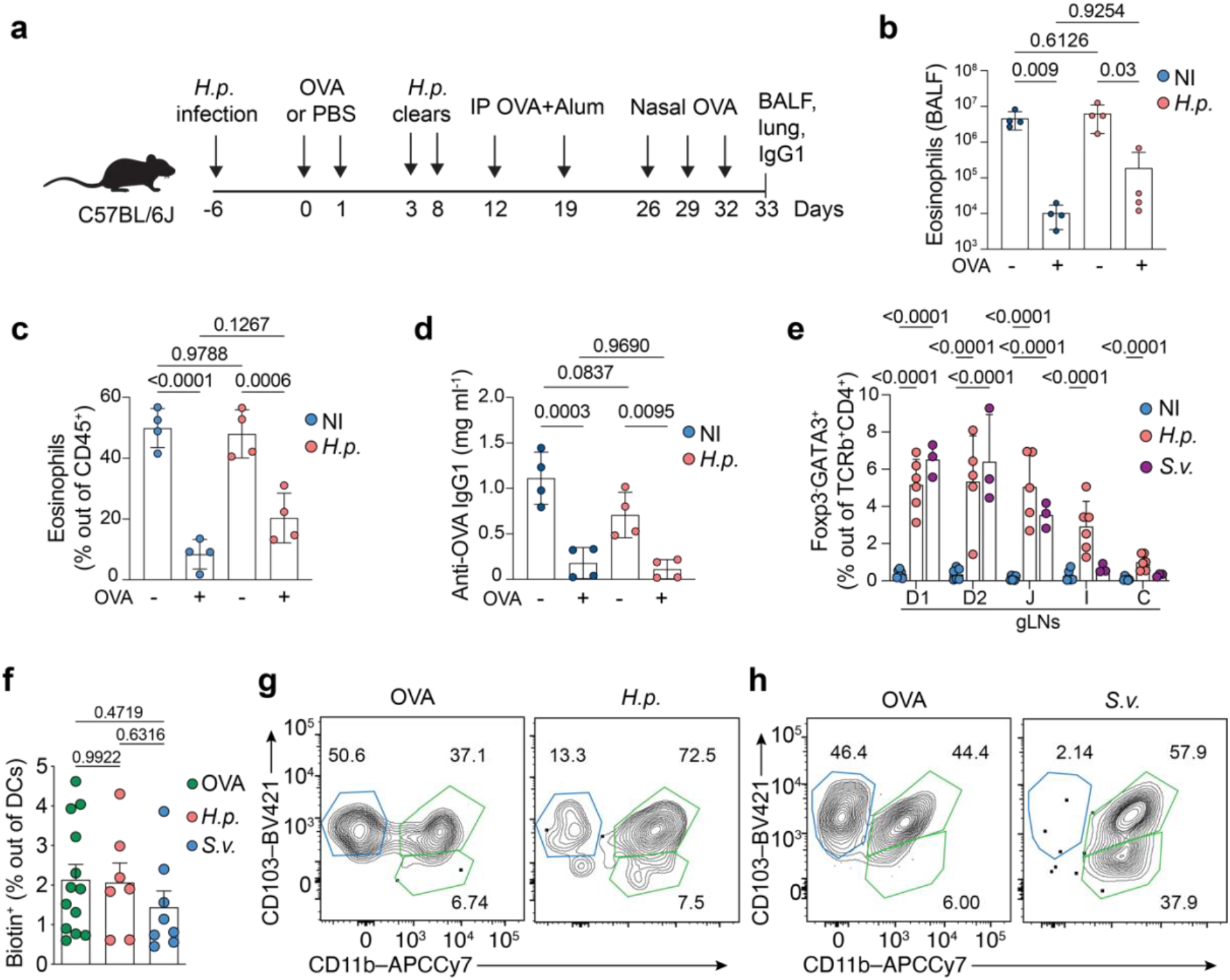
Supplementary Figure 3. *Heligmosomoides polygyrus* infection does not break oral tolerance. **(a-d)** Mice were infected with *Heligmosomoides polygyrus (H.p)* or not during antigen feeding (+OVA groups), or no feeding (−OVA groups), 18 days before first immunization with OVA–alum. **(a)** Scheme of oral tolerance experimental set up in *H.p*. - infected mice. **(b)** Total eosinophils in BALF, **(c)** Percentage of eosinophils among CD45^+^ cells in lung tissue and **(d)** OVA-specific IgG1 levels in serum (n = 4 mice per group, representative of two independent experiments). **(e)** Non-infected (NI), *Strongyloides. venezuelensis (S.v)* or *H.p* infected C57BL/6 CD45.2 mice were adoptively transferred with 1 × 10^6^ naive CD45.1^+^ CD4^+^ OT-II T cells and analyzed 64 h after cell transfer. Two doses of intragastric OVA were given at 48 h and 24 h prior analysis. Percentage of GATA3^+^ cells among CD45.2 TCRβ^+^CD4^+^ cells in gLNs (n = 3 mice per group, pool of two independent experiments). (**f-h**) CD45.2 *Cd40*^G5/G5^ mice were infected with *S.v*. or *H.p*. 5 days prior adoptively transfer of 1 × 10^6^ naive CD45.1 CD4^+^*Cd40lg*^SrtA/Y^ OT-II T cells. Animals received 1 dose of intragastric OVA and cell-cell interaction was revealed by LIPSTIC protocol 24 h later. Non-infected mice (OVA) were used as control. **(f)** Percentage of labeled DCs in the D-gLNs of OVA, *H.p.-*or *S.v*-infected mice (n = 2, 4 mice per group, pool of three independent experiments). Representative flow plots showing percentage of biotin^+^cDC1 (blue) and cDC2 (green) in D-gLNs of **(g)** *H.p* or **(h)** *S.v* infected mice. Quantification of data in Fig. 3c, d. *P*-values were calculated by one-way or two-way ANOVA.

**Supplementary Figure 4.**
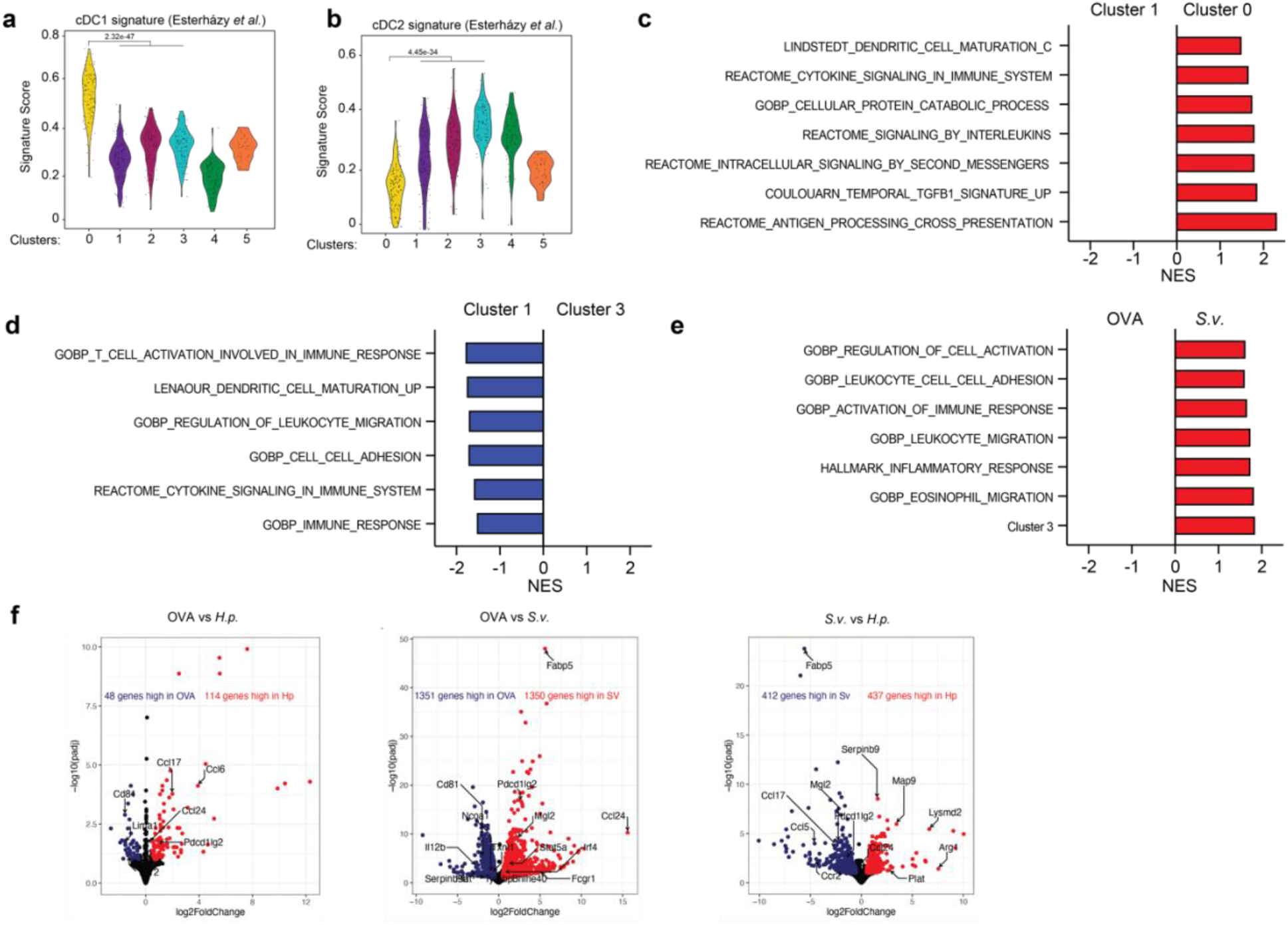
Transcriptomics of D-gLNs and duodenum lamina propria DCs from *Heligmosomoides polygyrus* or *Strongyloides venezuelensis* infected mice. Expression of **(a)** cDC1 and **(b)** cDC2 gene expression signatures, obtained from the literature^3^, in transcriptional Clusters as defined in Fig. 4a. Differentially expressed gene signatures of D-gLNs DCs between **(c)** Cluster 0 *vs* Cluster 1 or **(d)** Cluster 1 *vs* Cluster 3. **(e)** Differentially expressed gene signatures of duodenum lamina propria DCs between *S.v. vs* OVA. **(f)** Volcano plots depicting genes that are differentially expressed between OVA *vs H.p.,* OVA *vs S.v*. and *H.p. vs S.v. P*-values were calculated using Wilcoxon signed-rank test or by fgsea and corrected with FDR.

